# Identification of potent biparatopic antibodies targeting FGFR2 fusion driven cholangiocarcinoma

**DOI:** 10.1101/2024.09.16.613045

**Authors:** Saireudee Chaturantabut, Sydney Oliver, Dennie T. Frederick, Jiwan Kim, Foxy P. Robinson, Alessandro Sinopoli, Tian-Yu Song, Diego J. Rodriguez, Liang Chang, Devishi Kesar, Yao He, Meilani Ching, Ruvimbo Dzvurumi, Adel Atari, Yuen-Yi Tseng, Nabeel Bardeesy, William R. Sellers

## Abstract

Translocations involving FGFR2 gene fusions are common in cholangiocarcinoma and predict response to FGFR kinase inhibitors. However, the rate and durability of response are limited due to the emergence of resistance, typically involving acquired FGFR2 kinase domain mutations, and to sub-optimal dosing, relating to drug adverse effects. Here, we report the development of biparatopic antibodies targeting the FGFR2 extracellular domain (ECD), as candidate therapeutics. Biparatopic antibodies can overcome drawbacks of standard bivalent monoparatopic antibodies, which often show poor inhibitory or even agonist activity against oncogenic receptors. We show that oncogenic transformation by FGFR2 fusions requires an intact ECD. Moreover, by systematically generating biparatopic antibodies that target distinct epitope pairs along the FGFR2 ECD, we identified antibodies that effectively block signaling and malignant growth driven by FGFR2-fusions. Importantly, these antibodies demonstrate efficacy in vivo, synergy with FGFR inhibitors, and activity against FGFR2 fusions harboring kinase domain mutations. Thus, biparatopic antibodies may serve as new treatment options for patients with FGFR2-altered cholangiocarcinoma.

**Summary:** We identify biparatopic FGFR2 antibodies that are effective against FGFR2 fusion driven cholangiocarcinoma.

## INTRODUCTION

FGFR2 fusions are found across a variety of cancer types including in 10-15% of primary intrahepatic cholangiocarcinoma (ICC), a tumor of the bile duct epithelium (*1, 2*). While three FGFR1-3/4 inhibitors are approved for the treatment of ICC(*3*), positive trial results are tempered by a short duration of disease control (<9 months) and limited overall response rates (18-42%)(*4*). Major challenges of approved FGFR inhibitors include on-target, off-tumor adverse effects and the emergence of secondary resistance mutations, particularly V565 gatekeeper mutations(*3*). On-target hyperphosphatemia, attributable to the essential role of FGFR1 in phosphate homeostasis, limits the optimal therapeutic dosing of FGFR1-3 inhibitors(*5*). While the recently developed FGFR2 selective kinase inhibitor, RLY-4008, has shown clinical promise with increased response rates, its benefits are not durable. Consequently, although FGFR2-fusion-positive ICCs exhibit a strong and sustained dependence on FGFR2 signaling, targeting the pathway with kinase inhibitors alone is insufficient to achieve the desired therapeutic benefit.

Therapeutic antibodies against the extracellular domain (ECD) of FGFR2 could serve as complementary treatment modalities to FGFR kinase inhibitors, offering the potential for high specificity against FGFR2 and retaining efficacy in the context of kinase domain mutations. Importantly, the ECD, comprising ligand binding domains D2 and D3 and the autoinhibitory D1 domain, is retained in all cases of intracellular fusion events. Thus, the FGFR2 ECD may be amenable to antibody-mediated targeting, although there are key questions and hurdles to address to ensure optimal therapeutic development.

One such question is the uncertainty of whether ligand activation contributes to the transforming capacity of FGFR2 fusions, which has important implications for antibody design. In this regard, antibodies to receptor tyrosine kinases (RTKs) can potentially function by blocking signaling as well as through antibody-dependent cellular cytotoxicity (ADCC) or through cytotoxic payloads(*6–8*). However, traditional bivalent antibodies against receptor tyrosine kinases (RTK) are often marginally effective inhibitors of signaling and instead act through the latter mechanisms (*6–8*). Indeed, of the current monoclonal antibodies approved in cancer indications, less than 10% exhibit classical signaling pathway blockade, with over 60% exerting immune effector functions and over 25% classified as antibody-drug conjugates (ADCs)(*9*). Furthermore, the bivalent monoparatopic nature of traditional antibodies (*10*), can lead to agonistic activity due to their intrinsic tendency to crosslink, dimerize, and induce receptor activation(*11–13*). These data suggest that improvements in the activity of traditional monoparatopic bivalent antibodies could lead to more effective therapeutic antibodies. As a result, alternative antibody modalities have been explored.

In the present study we explored optimal approaches to antibody-mediated targeting of FGFR2 fusions. First, we defined the contributions of the FGFR2 ECD to transformation by FGFR2 fusion alleles. Secondly, we developed biparatopic antibodies as an improved strategy to targeting the FGFR2 ECD. Biparatopic antibodies, which recognize two distinct epitopes on the same protein, are a promising format which has been shown to generate highly potent antagonists (*14*). Here, by generating all 15 possible combinatorial heterodimeric biparatopic antibodies from 6 optimized monoparatopic antibodies that bind to distinct epitopes along the FGFR2 ECD, we identified two anti-FGFR2 biparatopic antibody candidates that are markedly superior to their parental bivalent antibodies in their potency against FGFR2 fusion driven cancers. Our study highlights the potential of biparatopic antibodies targeting FGFR2 as novel therapeutic agents.

## RESULTS

### The extracellular domain is necessary for full transformation by FGFR2 fusions

To ascertain the role of FGFR2-fusion ECDs, we developed BaF3 and NIH3T3 fibroblast cell lines expressing FGFR2 fusions: FGFR2-BICC1 (the most common fusion found in ICC), FGFR2-AHCYL1, and FGFR2-PHGDH. Expression of FGFR2 fusions resulted in IL-3-independent growth of BaF3 cells and transformation of NIH3T3 cells (Fig.1A and Fig.S1A); growth of these cells was robustly attenuated by the FGFR inhibitor (FGFRi) infigratinib (Fig.S1A). Transformation and proliferation of the FGFR2-fusion expressing lines were further enhanced by the FGFR2 ligand, FGF10 (Fig.1A, B). To determine the extent of receptor dimerization, we utilized NanoBiT assays that detect protein interactions by proximity-mediated luciferase fragment complementation(*15*) (Fig.1C). We validated expression of full-length FGFR2-WT and FGFR2-ACHYL1 coupled to the NanoBiT fragments, LgBiT and SmBiT (Fig.S1B) and assayed luminescent activity upon co-expression (Fig. S1C, D). Complementation-based luciferase activity of FGFR2 fusions were significantly higher than that of FGFR2-WT (Fig.1D), indicating ligand-independent dimerization. Nonetheless, addition of FGF10 significantly enhanced receptor dimerization of both FGFR2-WT and FGFR2-ACHYL1 (Fig.1D). These data indicate that the FGFR2-fusion ECD is functional and enhances receptor activation through ligand-mediated dimerization.

**Fig. 1.**
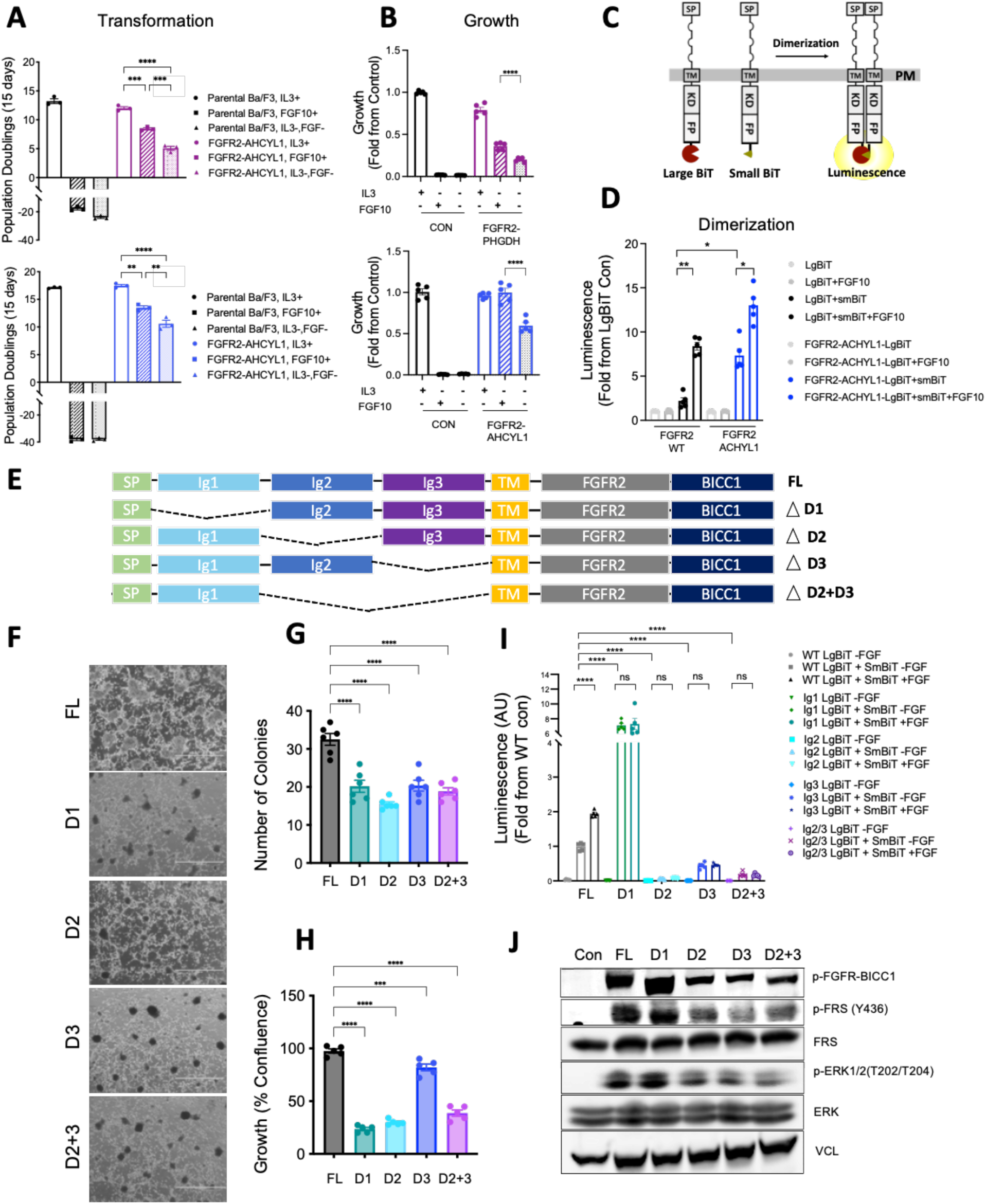
The extracellular domain is necessary for full transformation by FGFR2 fusions. A) Transformation assays showing cumulative population doublings in BaF3 cells expressing FGFR2-PGHDH (12 days) and FGFR2-ACHYL1 (15 days) with or without FGF10 or IL3 as indicated. B) Growth of BaF3 cells expressing FGFR2-PGHDH and FGFR2-ACHYL1 analyzed by CellTiter-Glo at 5 days post IL3 removal. C) Illustration of the dimerization assay using FGFR2-fusion NanoBiT constructs. Large BiT and Small BiT subunits are fused to the C-terminus of FGFR2 fusions. Upon receptor dimerization, SmBiT and LgBiT come together and produce luminescent light in the presence of substrate. SP: signal peptide, TM: transmembrane, KD: kinase domain, FP: fusion partner, PM: plasma membrane. D) HEK-293T cells expressing FGFR2-WT and FGFR2-ACHYL1 fused to LgBiT alone or fused to LgBiT and SmBiT were used to quantify the receptor dimerization in the presence or absence of FGF10. Shown is the fold increase over FGFR2-LgBiT activity alone. E) Illustration of FGFR2-BICC1 constructs with D1 (Ig1), D2 (Ig2), D3 (Ig3), or D2+D3 (Ig2+Ig3) deletions in the ECD. F) Representative images of focus formation assays of NIH-3T3 cells expressing FGFR2 WT or the indicated ECD deletion variants. G) Quantification of number of colonies from Fig. 1F. H) Growth of NIH3T3 cells overexpressing FL, D1, D2, D3, and D2+3 deleted FGFR2- BICC1 constructs as measured by Incucyte at 5 days post plating. I) Dimerization of FGFR2-BICC1 D1, D2, D3, or D2+D3 ECD deleted constructs in HEK- 293T cells compared to full-length FGFR2-BICC1. Fold change in luminescence over FGFR2-WT-LgBiT is shown. J) Immunoblotting of FGFR2 downstream pathway effectors in HEK-293 cells expressing FGFR2-BICC1 ECD deletion constructs. Data are mean ± SEM across three separate replicates. ns=not significant, *P < 0.05, **P < 0.01, ***P < 0.001, ****P < 0.0001 by One-way ANOVA multiple comparisons.

Next, we asked whether subdomains of the ECD were required for FGFR2-fusion dimerization and cell growth and transformation. To this end, we generated FGFR2 fusions with deletions of each of the subdomains, D1, D2, and D3 (Fig.1E). Since the D2 and D3 domains are necessary and sufficient for ligand binding, we also generated D2+3 deletion constructs. We next expressed each ECD deletion mutant in NIH3T3 cells lacking endogenous FGFR2 and performed colony formation and proliferation assays. Comparable expression of each construct was observed via immunoblotting (Fig.S1E). D1, D2, D3, and D2+3 deleted FGFR2 fusion expressing cells each showed significantly decreased growth (35-77% growth inhibition) and transformation capacity (36-50% reduction) compared to full length (FL) FGFR2 fusion expressing cells (Fig.1F, G, H). Specifically, deletion of D2 of the FGFR2 ECD had a pronounced impact on cell growth and transformation, suggesting that D2 may play a prominent role in the oncogenicity of FGFR2-BICC1. In all, the ECD is required for full transformation by FGFR2 fusions.

Signaling by wild type FGFR2 is initiated by the binding of FGF ligands to the D2 and D3 domains leading to FGFR dimerization and activation. To test whether these domains play comparable roles in the activation of FGFR2 fusions, we again utilized NanoBiT complementation and immunoblotting assays. The D2, D3, and D2+3 deleted FGFR2 fusions showed significantly impaired dimerization in the presence or absence of FGF10 ligand (Fig.1I). In keeping with the identification of D1 as the autoinhibitory domain of FGFR1 and FGFR3 (*16*), loss of the D1 domain enhanced receptor dimerization. Finally, we assessed the downstream pathway activation of the ECD deletion constructs by immunoblotting. Compared to the FL construct, expression of the D2, D3, and D2+3 deletion derivatives resulted in markedly impaired FGFR2 signaling, as reflected by reduced p-FGFR2, p-FRS2, and p-ERK, whereas deletion of D1 increased pathway activation, correlating with the observed increase in dimerization ((Fig.1I, J). Together, these data demonstrate that the FGFR2-fusion ECD is necessary for full oncogenic transformation of FGFR2 fusions. We further identify the D1 domain as an autoinhibitory domain, deletion of which activates ERK and diminishes viability through pathway activation, consistent with previous observations of activation dependent lethality we and others have observed in the setting of BRAF and NRAS mutations(*17, 18*).

### Development of candidate biparatopic antibodies directed against FGFR2

To determine whether biparatopic antibodies can disrupt the function of FGFR2 fusions, we identified and produced 6 optimized FGFR2 antibodies(*19–22*), including the parental antibody of bemarituzumab, an FGFR2 antibody with enhanced effector function undergoing phase III clinical testing(*23*). Available data suggested these antibodies likely bind to distinct epitopes in the ECD of FGFR2b, the primary isoform of FGFR2 fusions expressed in ICC(*3*). We compared and validated the reported binding epitopes and binding affinities, ascertaining FGFR2 binding by flow cytometry and Bio-Layer Interferometry (BLI) Octet analysis. Flow cytometric analysis confirmed high FGFR2 binding affinities of parental antibodies A-F, with dissociation constants (Kd) ranging from 0.15nM-32.79nM (Fig.2A). To validate their binding epitopes, NIH3T3 cells expressing FGFR2-fusion constructs with N-terminal deletions in D1, D2, D3, or D2+3 (Fig.1E) were analyzed by flow cytometry. The data showed that antibody A bound to all constructs, antibody B bound to all except the D1-deleted construct, antibodies C and D bound to all but the D2-deleted construct, and antibodies E and F bound to all except the D3-deleted construct (Fig.2B). These data suggest the following binding epitopes: antibody B (D1), antibodies C and D (D2), antibodies E and F (D3), and antibody A (outside the D1-3 domains, likely involving the N-terminus), consistent with prior reports(*20*). BLI-Octet epitope binning analysis by pairwise cross-competition corroborated our findings, showing antibodies A and B with unique binding epitopes while antibodies C, D and antibodies E, F pairs having overlapping epitopes (Fig.2C, D).

**Fig. 2.**
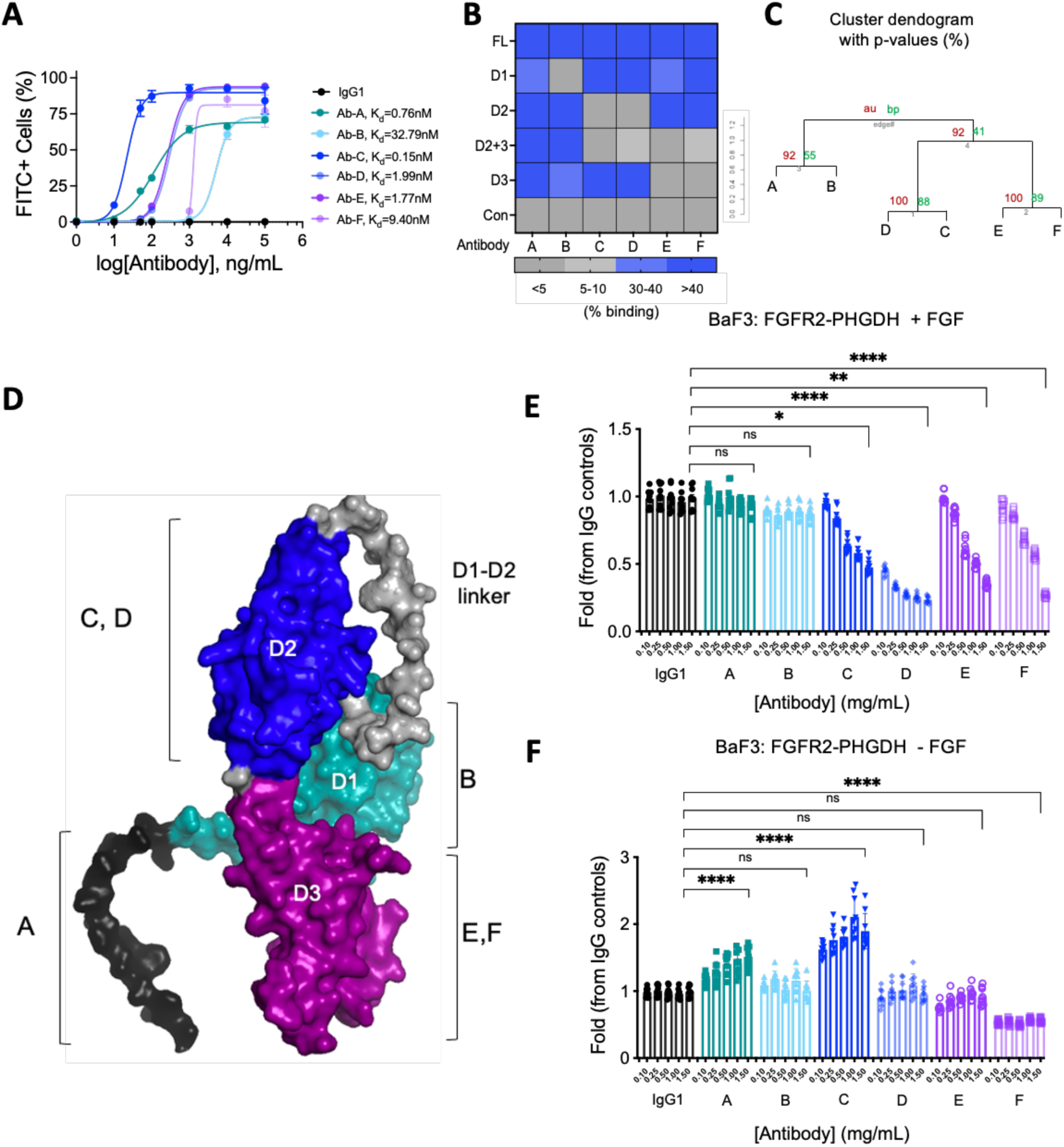
Development of candidate biparatopic antibodies directed against FGFR2 A) Binding affinities of the indicated anti-FGFR2 antibodies via flow cytometry. B) Flow cytometry analysis using anti-hIgG1-FITC secondary antibody to detect FGFR2 parental antibodies A-F. Binding epitopes of parental antibodies A-F along the FGFR2 ECD were identified using full-length, D1, D2, D3, and D2+3 deleted FGFR2-BICC1 overexpressing NIH3T3 cell lines shown in Fig.1. C) Epitope binning through cross competition assay. BLI-Octet Epitope clustering diagrams showing cluster dendrogram with au (approximately unbiased) p-values and bp (bootstrap probability) value (%). Distance represents correlations and cluster method is average. D) Alpha-fold predicted structure of FGFR2 ECD showing D1, D2, D3 and D1-D2 flexible linker as well as 6 FGFR2 parental antibody binding epitopes A-F. E-F) Viability of FGFR2-PHGDH overexpressing BaF3 cells upon treatment with increasing concentrations of antibody A-F in the presence or absence of FGF10 ligand. Data are mean ± SEM across seven separate replicates. ns=not significant, *P < 0.05, **P < 0.01, ***P < 0.001, ****P < 0.0001 by One-way ANOVA multiple comparisons.

To determine whether targeting FGFR2-fusion ECDs with anti-FGFR2 antibodies impairs their oncogenic activity, we treated BaF3 cells expressing FGFR2-PHGDH with each parental FGFR2 antibody. Antibodies against the ligand-binding ECD (antibodies C, D, E, and F) inhibited FGF-stimulated growth (Fig.2E). These results support our hypothesis that ligand-bound ECD augments FGFR2-fusion activity and that the ECD is necessary for FGFR2 fusion driven growth. In the ligand-independent setting, only antibody F inhibited FGFR2-PHGDH driven BaF3 cell growth (Fig.2F). Antibodies B, D, and E had marginal impact on cellular growth in this setting, while antibodies A and C exhibited agonistic activity and promoted ligand-independent growth (Fig.2F). Consistent with its agonist activity, antibody C increased dimerization of FGFR2-ACHYL1 and FGFR2- BICC1 in our FGFR2 NanoBiT dimerization assay (Fig.S2A). Bivalent, monoclonal antibodies against the MET receptor are well known to agonize and dimerize the receptors(*10*). Thus, the growth-promoting effects observed with anti-FGFR2 antibodies A and C in the absence of ligand could be similarly due to their bivalent nature. In addition, the differential behavior of antibodies C and D suggests that they bind to distinct epitopes within the D2 domain.

We next asked whether FGFR2 biparatopic antibodies might have enhanced potency and avoiding ligand-independent receptor activation. To this end, we used controlled Fab-arm exchange to generate full IgG1 FGFR2 antibodies that simultaneously bind two different epitopes on the FGFR2 ECD(*24*). Here, complementary IgG Fc mutations allow preferential heterodimer formation between distinct IgG-formatted antibodies while maintaining heavy and light chain pairing. We produced each of the 6 parental antibodies with the reciprocal mutations, allowing us to create 15 novel biparatopics from all possible pairwise combinations (Fig.3A, B). We validated the purity of the biparatopic antibodies via mass spectrometry (exemplified in Fig.S3A, B). Each biparatopic antibody showed >95% purity with little to no residual from either parental antibody. In all, we validated the binding affinities as well as binding epitopes of the 6 parental antibodies and generated 15 biparatopic antibodies for further characterization.

**Fig. 3.**
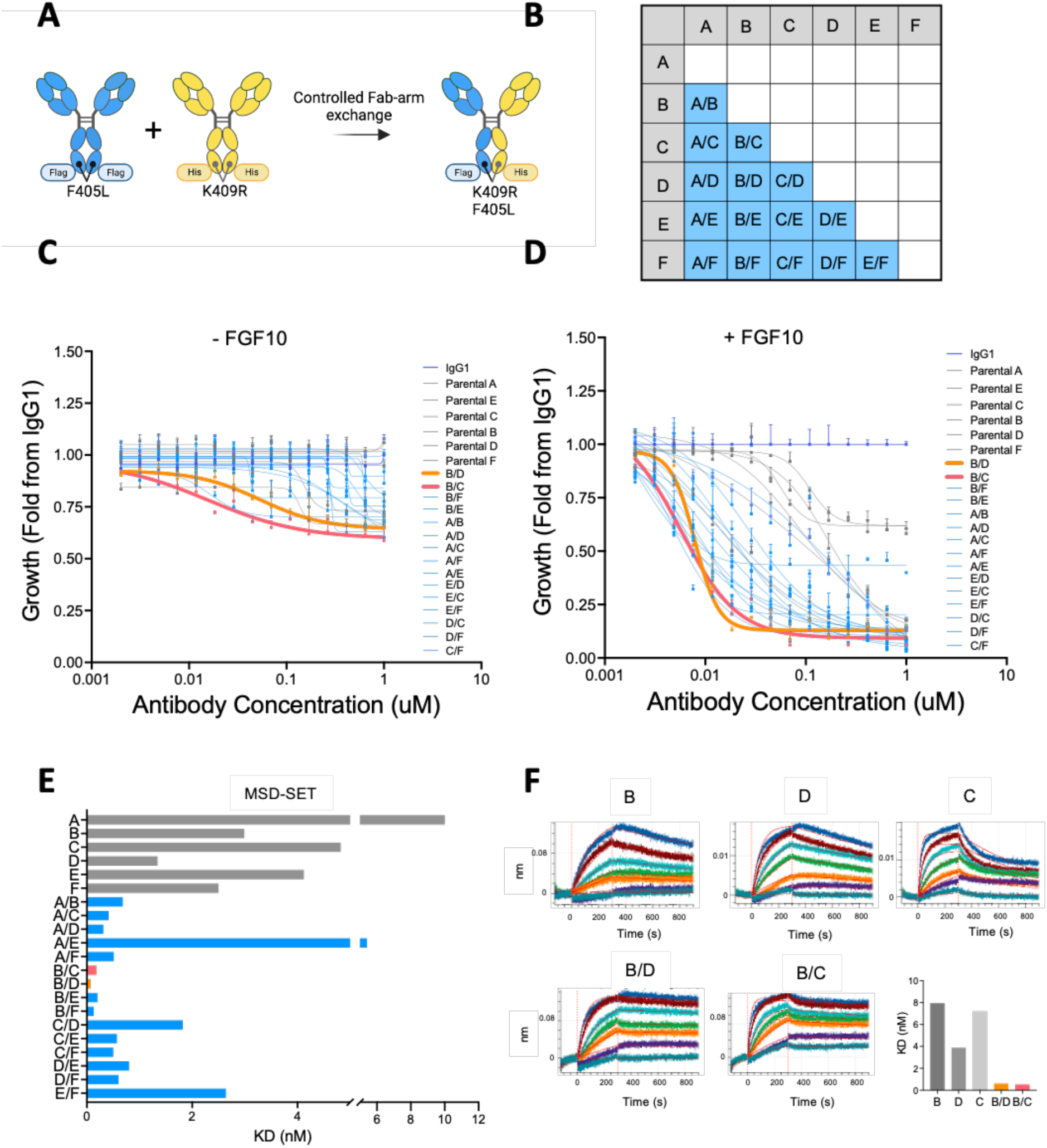
Identification of potent tumor growth-inhibiting biparatopic antibodies via unbiased screening A) Illustrations showing strategy for biparatopic antibody generation. B) A diagram showing all 15 possible biparatopic antibody pairs that were generated from 6 parental antibodies A-F. C-D) Viability of FGFR2-ACHYL1 overexpressing BaF3 cells upon treatment with IgG1, biparatopic antibodies, and their parental antibodies in the absence (C) and presence of FGF10 (D) E) Binding affinities (KD, nM) of parental antibodies (gray) compared to biparatopic antibodies (blue) from MSD-SET assay. Biparatopic antibodies bpAb-B/D and bpAb-B/C with binding affinities KD of 0.07nM (orange bar) and 0.18nM (pink bar) respectively. F) Representative binding sensorgrams illustrating the binding kinetics between FGFR2 ECD and antibody B, D, C or biparatopic antibody bpAb-B/C and bpAb-B/D via BLI- Octet. KD values were determined based on the association and dissociation curves.

### Unbiased screening identifies potent, tumor growth inhibiting biparatopic antibodies

To determine the potency of each biparatopic antibody in inhibiting tumor cell growth, we assessed antiproliferative activity in FGFR2 fusion driven BaF3 cells with or without addition of ligand. Of the 15 biparatopic antibodies tested, 7 (46%) and 11 (73%) outperformed parental antibodies at inhibiting growth of FGFR2-ACHYL1 driven BaF3 cells in the absence or presence of FGF10 ligand, respectively (Fig 3C, D). A second BaF3 model driven by an FGFR2-PHGDH fusion yielded similar results (Fig S3C, D). Notably, bpAb-B/C and bpAb-B/D were the most potent of the 21 parental and biparatopic antibodies in the viability assays. Importantly, the efficacy of pairwise mixtures of the parental antibodies differed from and did not predict the potency of their respective biparatopic antibodies (Fig.S3E, F), suggesting that novel modes of action are enabled by the biparatopic antibody generation.

We next determined the affinity of the biparatopic antibodies for FGFR2. Using the MSD- SET assay, we found that 80% (12 out of 15) of biparatopic antibodies, including bpAb- B/C and bpAb-B/D, had significant improvements (>10 fold) in FGFR2 binding affinities as compared to their parental antibodies (Fig.3E). The remaining 3 biparatopic antibodies with lower affinities had binding epitopes either within the same ECD subdomain (D2 for bpAb-C/D; D3 for bpAb-E/F) or on subdomains that are the furthest apart (D1 and D3 for bpAb-A/E). These data suggest that the geometry of binding between antibodies and their epitopes plays an important role in achieving high affinity binding. We then examined the kinetics of antibody association and dissociation using BLI-Octet analysis. In addition to their enhanced binding avidity, antibodies bpAb-B/C and bpAb-B/D also exhibited significantly slower off-rates compared to their parental antibodies B, C, and D (Fig.3F). Both bpAb-B/C and bpAb-B/D contain binding arms against epitope B, a flexible autoinhibitory extracellular domain (ECD) D1 (Fig.2D). Together, our data demonstrate that the majority of biparatopic antibodies against combinations of selected epitopes on the FGFR2 ECD have enhanced efficacy and avidity compared to their parental antibodies. Based on their improved potency and avidity, we selected bpAb-B/C and bpAb-B/D for further characterization.

### Biparatopic antibodies show superior inhibition of growth and transformation of FGFR2 fusion driven cholangiocarcinoma cell lines

We investigated the impact of biparatopic FGFR2 antibody candidates bpAb-B/C and bpAb-B/D on two patient-derived models of FGFR2 fusion+ ICC, ICC13-7 (FGFR inhibitor-sensitive) and ICC21 (partially sensitive) (*25*). They endogenously express the FGFR2-OPTN and FGFR2-CBX5 fusions, respectively. Correlating with their activity in FGFR2 -fusion expressing BaF3 cells, bpAb-B/C and bpAb-B/D have enhanced efficacy at inhibiting growth of ICC13-7 and ICC21 cells in the absence (Fig.4A, C) and, even greater, in the presence (Fig.4B, C) of FGF10 compared to the parental antibodies.

**Fig. 4.**
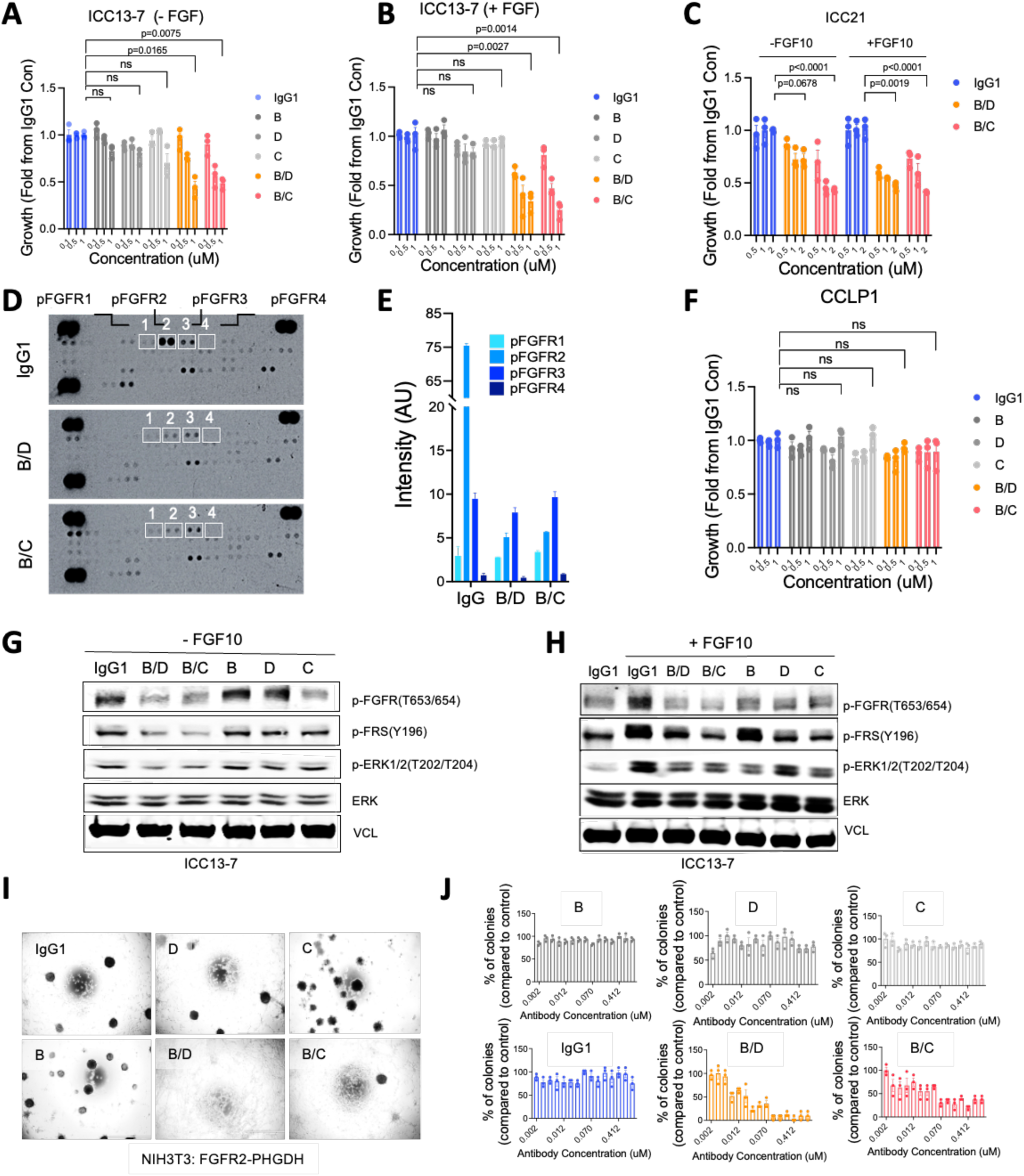
Biparatopic antibodies show superior inhibition of growth and transformation of a FGFR2 fusion-driven cholangiocarcinoma cell line. A-C) Viability of cholangiocarcinoma cell line ICC13-7 or ICC21 upon treatment with biparatopic antibodies bpAb-B/C, bpAb-B/D, parental antibodies B, D, C or IgG1 isotype in the absence (A, C) or presence (B, C) of FGF10. ICC13-7 or ICC21 cells were grown in RPMI media containing 1% FBS and were treated every 3 days. Growth was measured by incucyte at 14 days post seeding. D-E) Proteome profiler human phospho-kinase array demonstrating levels of 43 phosphorylated human kinases in NIH3T3 cells overexpressing FGFR2-PHGDH treated with IgG1, bpAb-B/C, or bpAb-B/D. bpAb-B/C and bpAb-B/D (D). Levels of p-FGFR1, p- FGFR2, p-FGFR2, and p-FGFR4 (white boxes) were analyzed and quantified (E) using ImageJ. F) Viability of CCLP-1 cells upon treatment with biparatopic antibodies bpAb-B/C, bpAb- B/D, parental antibodies B, D, C or IgG1 isotype control. G-H) Immunoblot of ICC13-7 cells upon 5hrs after treatments with bpAb-B/C, or bpAb- B/D compared to the parental antibodies B, D, C in the absence (G) or presence (H) of FGF10 ligand. I-J) Representative images of focus formation assays of FGFR2-PHGDH expressing NIH3T3 cells upon treatments with parental antibodies B, D, C, biparatopic antibodies bpAb-B/C and bpAb-B/D or IgG1 (I) as quantified by the number of colonies (J). Data are mean ± SEM across three separate replicates. ns=not significant, *P < 0.05, **P < 0.01, ***P < 0.001, ****P < 0.0001 by One-way ANOVA multiple comparisons.

To investigate whether cell growth inhibition caused by bpAb-B/C and bpAb-B/D were specific to inhibition of FGFR2 rather than other FGFRs, extracts from NIH3T3 cells expressing FGFR2-PHGDH were profiled using a phospho-RTK array. We found that bpAb-B/C and bpAb-B/D specifically inhibited phosphorylation of FGFR2 but not of FGFR1 or FGFR3 (Figs. 4D, E; minimal FGFR4 phosphorylation was detected in these cells). We also tested FGFR2 specificity using the CCLP-1 ICC cell line, which lacks an FGFR2 fusion and is driven by FGFR1 and FGF20 overexpression(*26*). Both bpAb-B/C and bpAb-B/D treatments had no significant impact on CCLP-1 cell viability, whereas the IC50 for FGFR1-3 inhibitor futibatinib is <1.5 nM (*26*) (Fig.4F). Thus, bpAb-B/C and bpAb- B/D inhibit FGFR2 with high specificity.

We next examined the effects of bpAb-B/C and bpAb-B/D on FGFR2-fusion mediated signaling. Both bpAb-B/C and bpAb-B/D significantly decreased p-FGFR2, p-FRS2, and p-ERK as compared to their parental antibodies B, C, or D in a ligand-independent setting (Fig.4G, S4A); additionally, bpAb-B/C and bpAb-B/D blocked FGF10-induced phosphorylation of FGFR, FRS2, and ERK (Fig.4H, S4A). Similarly, bpAb-B/C and bpAb- B/D impaired downstream signaling in NIH3T3 cells expressing FGFR2-PHGDH (Fig.S4B). Thus, bpAb-B/C and bpAb-B/D specifically inhibit downstream signaling by constitutively active FGFR2-fusion proteins.

We next assessed the ability of bpAb-B/C and bpAb-B/D to inhibit FGFR2-fusion driven oncogenic activity via focus formation assays using FGFR2-PHGDH transformed NIH3T3 fibroblasts (Fig.4I). Cells treated with bpAb-B/C and bpAb-B/D showed a dose-dependent decrease in transformation capacity (reduction in colony formation), whereas the parental antibodies and IgG1 treated control had no effect (Fig.4J). Collectively, these results highlight the specificity of the biparatopic antibodies towards FGFR2 and the significant improvement in the potency of FGFR2 inhibition when compared to bivalent monotopic antibodies.

### Biparatopic antibodies show superior in vivo anti-tumor activity compared to the parental antibodies

We next tested the in vivo efficacy of bpAb-B/C and bpAb-B/D and their parental antibodies against subcutaneous tumors formed by FGFR2-PHGDH transformed BaF3 cells in SCID mice. At a tumor size of ∼250mm^3^, mice were randomized into 10 groups with 10 mice per treatment group. The antibodies were administered via intravenous tail vein injections twice per week. Both bpAb-B/C and bpAb-B/D biparatopic antibodies potently suppressed tumor growth at 5, 15, and 25mg/kg doses, whereas the parental antibodies (administered at 15 mg/kg) showed no anti-tumor activity (Fig.5A, B). Pharmacokinetics analysis by ELISA demonstrated dose-proportional increases in the plasma concentration of the biparatopic antibodies, and furthermore, significantly longer half-life compared to small molecule inhibitors, consistent with their larger size (*27, 28*) (Fig.S5A, B).

**Fig. 5.**
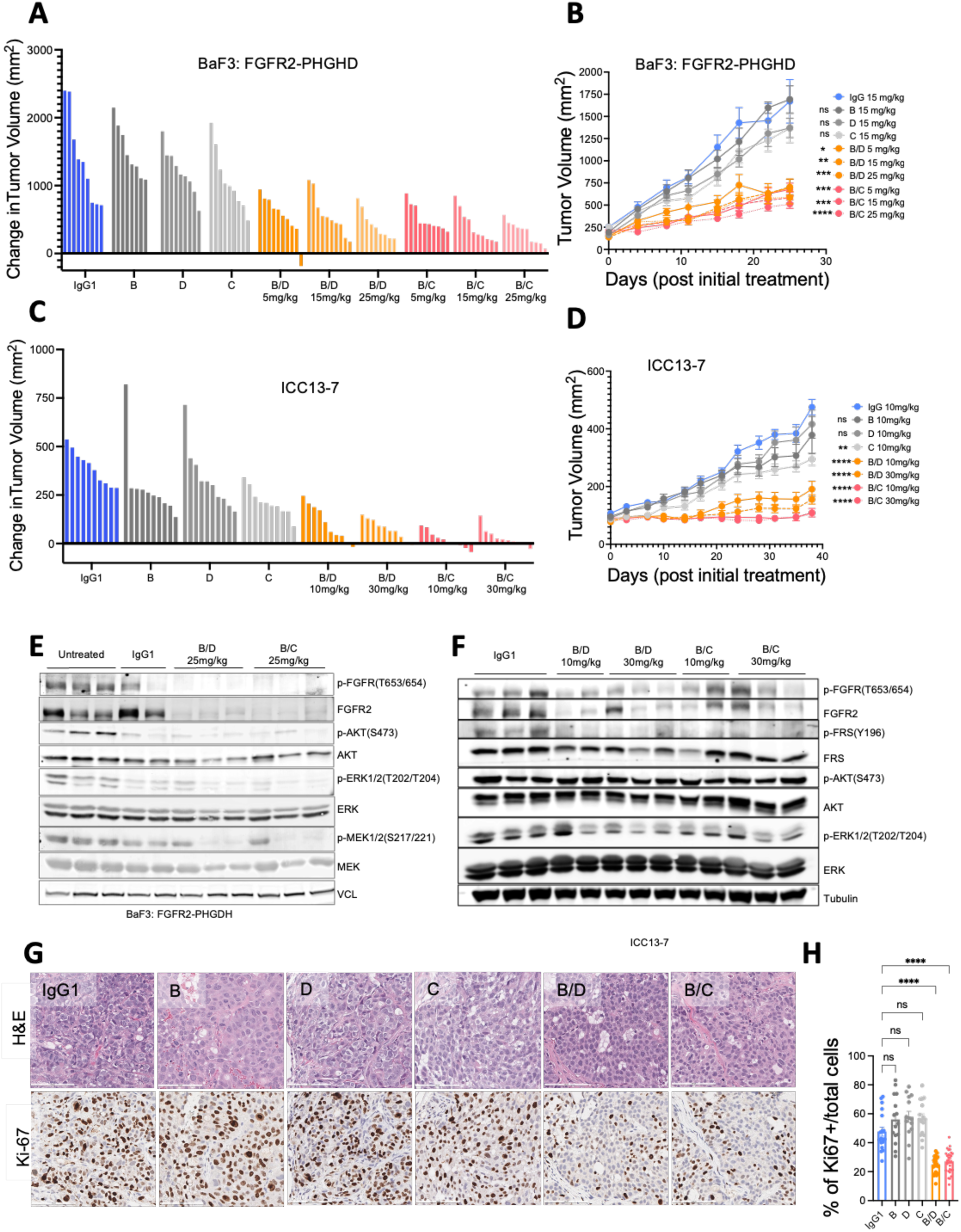
Biparatopic antibodies show superior in vivo anti-tumor activity compared to the parental antibodies. A-D) BALB/c scid mice harboring BaF3 cells overexpressing FGFR2-PHGDH (A, B) or ICC13-7 (C, D) subcutaneous xenografts were treated with antibodies when the tumor reached ∼ 250mm^3^ (A, B) or 150mm^3^ (C, D). Mice (n=10 per group) were administered twice weekly with IgG1, parental antibodies B, D, C, at indicated dosage via IV injections. Results are represented in the waterfall plot illustrating changes in tumor volume at day 25 (A, B) or day 38 (C, D) post initial treatment (A, C) and as geometric mean of tumor volumes ± SEM every 3-4 days from day 0-day 25 post initial treatment (B, D). Data are mean ± SEM across ten mice. ns=not significant, *P < 0.05, **P < 0.01, ***P < 0.001, ****P < 0.0001 by Friedman’s ANOVA multiple comparisons. E) Immunoblot analysis of FGFR2-PHGDH overexpressing BaF3 cells xenograft tumors. Tumors were harvested 5 hrs after the final round of bpAb-B/C, bpAb-B/D, or IgG1 administration. F) Immunoblot analysis of ICC13-7 xenograft tumors collected 5 hrs after the final round of antibody administration on day 38. G) Representative images of hematoxylin and eosin stains (H&E) and immunohistochemistry (IHC) staining for proliferation marker Ki-67 in ICC13-7 xenograft tumor samples on the final day of treatment. Scale bars, 100um. H) Quantification of % number of Ki-67 positive nuclei normalized to the total number of nuclei (nuclei counterstain). Data are from 2 biological replicates per treatment group with at least 14 representative images for analysis per group. Data are mean ± SEM. ns=not significant, *P < 0.05, **P < 0.01, ***P < 0.001, ****P < 0.0001 by One-way ANOVA multiple comparisons.

The biparatopic antibodies also showed prominent i*n vivo* efficacy against xenograft tumors formed by the patient-derived, ICC13-7 cholangiocarcinoma model. While the parental antibodies had only marginal effects on tumor growth, the biparatopics were highly effective at both 10 and 30 mg/kg dose concentrations. Notably, bpAb-B/C showed greatest potency, resulting in tumor stasis at 38 days post-treatment (Fig.5C, D), comparable to the efficacies of clinically used FGFR inhibitors(*25, 29*). Importantly, bpAb- B/C and bpAb-B/D treatment in both in vivo models led to a significant decrease in total FGFR2 levels and reductions in p-FGFR, p-FRS and p-ERK compared to IgG1 control (Fig.5E, F). By contrast, the parental antibodies showed limited effect on total FGFR2 levels or on downstream signaling (Fig.S5C). Consistent with the tumor growth inhibition data, bpAb-B/C and bpAb-B/D markedly decreased tumor cell proliferation (Ki-67 staining) compared to parental antibodies or IgG1 control (Fig 5G, H). None of the antibody treatment affected mouse body weight (Fig.S5D, E). Assessment of antibody tumor distribution by IHC staining showed that bpAb-B/C and bpAb-B/D localized to the cell membrane and exhibited diffuse staining throughout ICC13-7 xenografts (Fig.S5F). Thus, although high affinity antibodies have been suggested to be restricted in their ability to penetrate tumors(*30*), these data demonstrate that biparatopic antibodies penetrate tumor effectively. Together, these results demonstrate that bpAb-B/C and bpAb-B/D have significantly improved anti-tumor activity compared to their parental antibodies in vivo.

### Biparatopic antibodies promote receptor internalization and lysosomal degradation

Based on the observation that bpAb-B/C and bpAb-B/D treatments significantly reduced the total levels of FGFR2 in vivo, we explored the impact of the biparatopic antibodies on receptor internalization and degradation. To determine whether bpAb-B/C and bpAb-B/D promote FGFR2-fusion internalization, we performed an internalization assay. FGFR2- PHGDH expressing BaF3 cells were treated with bpAb-B/C, bpAb-B/D, or IgG control before being transferred to 4°C to block or 37°C to induce internalization, and surface FGFR2 was analyzed by flow cytometry (Fig.6A, B). Cells treated with bpAb-B/C and bpAb-B/D showed increased internalization from 60-960 minutes (from ∼6% to 80% shift in surface FGFR2) (Fig.6B). The internalization assay was repeated in ICC13-7 cells treated with bpAb-B/C, bpAb-B/D, respective parental antibodies, or IgG control. ICC13- 7 cells treated with bpAb-B/C and bpAb-B/D had a significant decrease in surface FGFR2 compared to cells treated with parental antibodies B, C, or D or IgG1, suggesting that bpAb-B/C and bpAb-B/D enhanced FGFR2 receptor internalization (Fig.6C). To investigate whether bpAb-B/C and bpAb-B/D treatments induce lysosome-mediated receptor trafficking, we labeled Fc-containing biparatopic and parental antibodies with a Fab fragment conjugated to a pH-sensitive fluorophore(*31*) and assessed lysosome-mediated induction of fluorescence in FGFR2-PHDGH, FGFR2-ACHYL1, and FGFR2-BICC1 expressing NIH3T3 cells (Fig.6D). Treatment with bpAb-B/C and bpAb-B/D resulted in significant increases in the fluorescent signal compared to the parental antibodies (Fig.6E-H). Moreover, labelling of lysosomes with lysotracker (green) and biparatopic antibodies with Fab-Fluor (red) demonstrated colocalization of the two signals, confirming the presence of the antibodies in the lysosomes (Fig.S6A). Consistent with results in FGFR2 fusion expressing NIH3T3 cells, treatment of the ICC13-7 cholangiocarcinoma cell line with bpAb-B/C and bpAb-B/D led to increases in fluorescent signals compared to parental antibodies (Fig.6I).

**Fig. 6.**
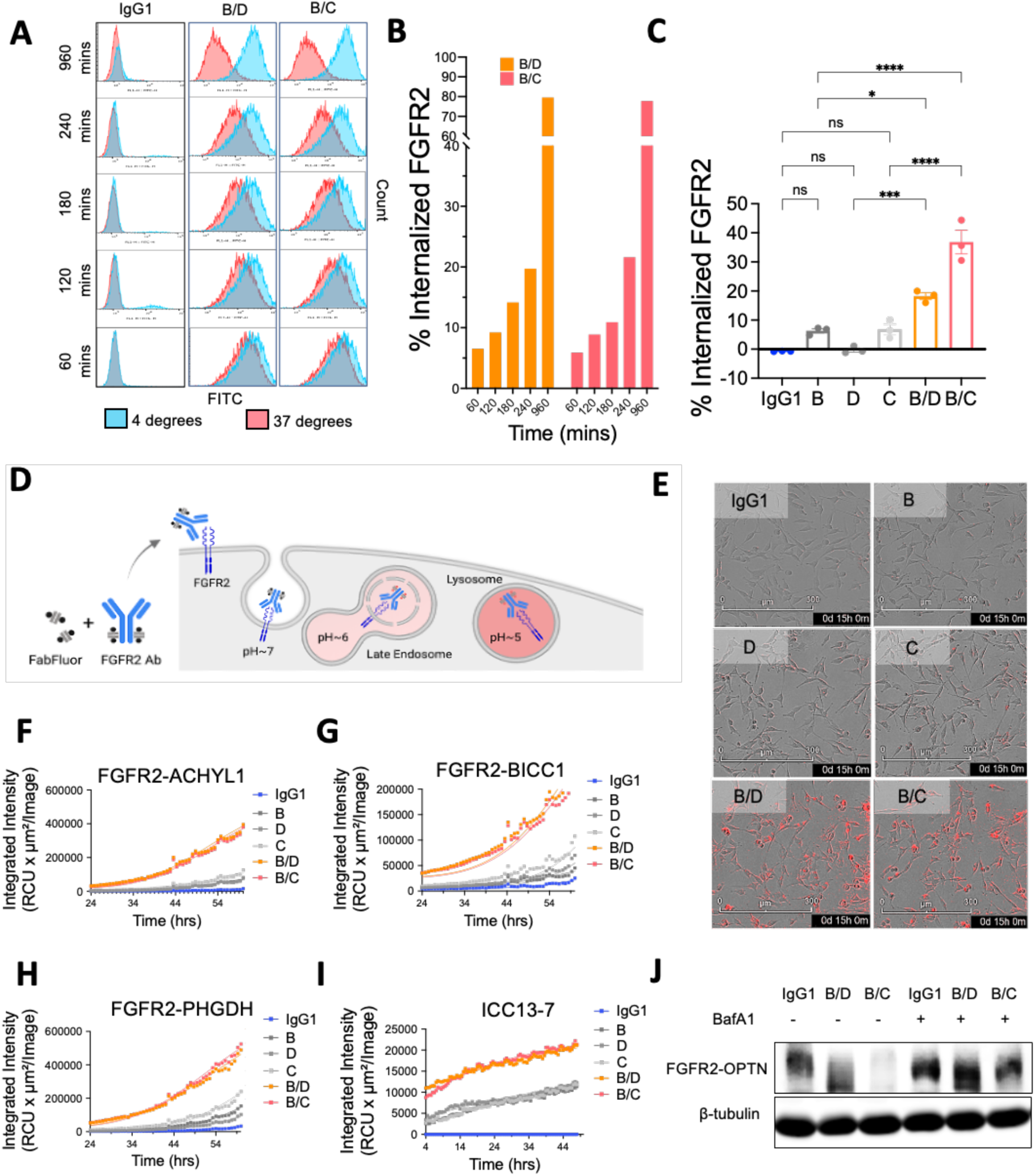
The biparatopic antibodies promote receptor internalization and lysosomal degradation. A) Flow cytometry histograms of surface FGFR2-PHGDH in BaF3 cells at 4 degrees Celsius (blue) and 37 degrees Celsius (red) upon treatment with bpAb-B/C or bpAb-B/D from 60-960 mins. B) Quantification of the histograms demonstrating the percentage of internalized FGFR2 at 60 mins, 120 mins, 180 mins, 240 mins, and 960 mins post bpAb-B/C or bpAb-B/D incubation. C) Quantification of histograms showing % internalized FGFR2 in ICC13-7 cell line at 4°C and 37°C after 5hrs of treatment with parental antibody B, D, C or biparatopic antibodies bpAb-B/C or bpAb-B/D. Data are mean ± SEM. ns=not significant, *P < 0.05, **P < 0.01, ***P < 0.001, ****P < 0.0001 by One-way ANOVA multiple comparisons. D) Illustrations of Fabfluor-pH antibody labeling assay. The pH sensitive dye-based system exploits the acidic environment of the lysosomes to quantify internalization of the labeled antibody. Fluorescent signals which indicate the internalization/degradation events were tracked using Incucyte. E) Representative images of detected fluorophore in NIH3T3 cells expressing FGFR2- PHGDH treated with parental antibody B, D, C or biparatopic antibody bpAb-B/C and bpAb-B/D at 15hr post incubation. F-H) Quantification of internalization/degradation signals in FGFR2-ACHYL1 (F), FGFR2-BICC1 (G), FGFR2-PHGDH (H) expressing NIH3T3 cells treated with parental antibodies B, D, C or biparatopic antibody bpAb-B/C and bpAb-B/D from 24 hrs post incubation. I) Quantification of internalization/degradation signals in ICC13-7 cells treated with parental antibodies B, D, C or biparatopic antibody bpAb-B/C and bpAb-B/D at 4 hrs post incubation. J) Immunoblot of ICC13-7 cells treated with IgG1, bpAb-B/C or bpAb-B/D antibodies alone or cotreated with bafilomycin A1 (BafA1) for 24 hrs. BafA1 was preincubated for 1hr prior to antibody treatments.

To investigate whether the observed increase in FGFR2 internalization and lysosomal trafficking mediate lysosomal degradation, we suppressed lysosome acidification and catabolism using the vacuolar-type H+-ATPase inhibitor bafilomycin A1 (BafA1). Bafilomycin treatment blocked bpAb-B/C- or bpAb-B/D-induced FGFR2-OPTN downregulation in ICC13-7 as compared to IgG1 treated control (Fig.6J). Together, these data demonstrate that bpAb-B/C and bpAb-B/D induce FGFR2-fusion internalization, trafficking, and lysosomal-mediated degradation to decrease FGFR2 fusion driven activity and growth. Notably, this mode of action induced by the biparatopic antibodies does not require co-targeting of lysosome-targeting receptors, membrane E3 ligases, or autophagy signaling molecules as seen in the development of LYTAC, AbTAC, or AUTAC systems(*32*).

### Biparatopic antibodies potentiate the efficacy of FGFR inhibitors

Given the specificity of FGFR2 antibodies and the potency of FGFR1-3 kinase inhibitors, combining two distinct treatment modalities might result in cooperativity specific to FGFR2 while sparing FGFR1 and 3, leading to more potent FGFR2 inhibition. To test whether bpAb-B/C and bpAb-B/D synergize with FGFRi, FGFR2-PHGDH expressing BaF3 cells were treated in a titration matrix of bpAb-B/C or bpAb-B/D in combinations with approved FGFRi infigratinib, futibatinib, and pemigatinib. The Bliss model was then applied to determine the degree of synergy (*33*). Bliss scores of 0-10 generally indicate additive interactions, while scores >10 demonstrate synergistic interactions. In the absence of FGF10, combination of bpAb-B/D with infigratinib, pemigatinib, or futibatinib as well as combination of bpAb-B/C with futibatinib or pemigatinib moderately enhanced growth inhibition (Fig.7A, B). Synergy between bpAb-B/C and infigratinib in a ligand-independent setting was striking, with a Bliss score of >20 (Fig.7B, C). In the presence of FGF10, co-treatments of bpAb-B/C or bpAb-B/D with infigratinib, futibatinib, and pemigatinib all enhanced growth suppression compared to treatment with single agents (Fig.7A-C). In accordance with the dose-response, all Bliss values were well above 10 in the ligand-dependent context (Fig.7C). These data highlight the potential of the biparatopic antibodies to boost the activity of FGFR inhibitors both in the presence and absence of ligand.

**Fig. 7.**
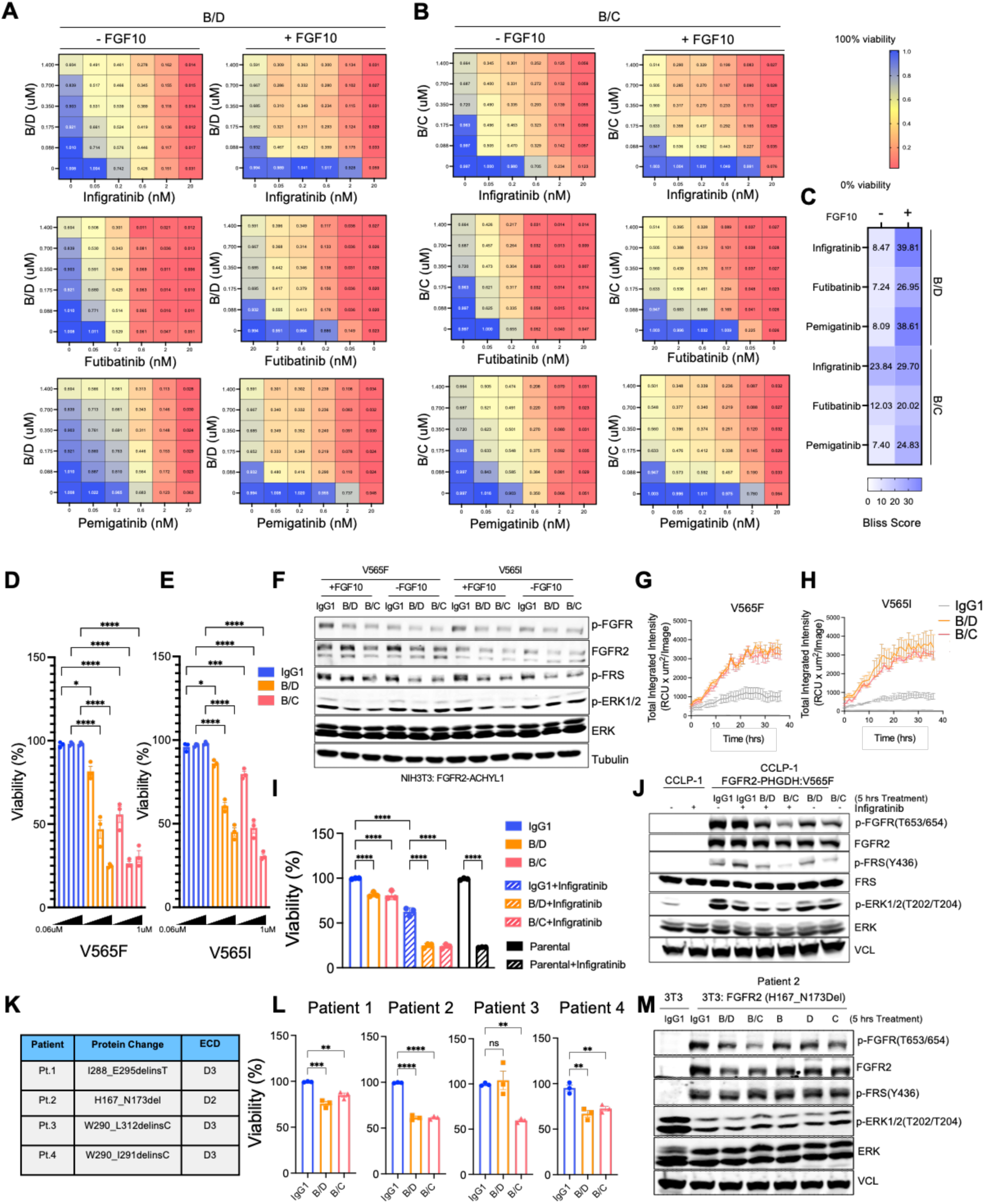
Combinations of biparatopic antibodies with FGFR inhibitors A) Biparatopic antibody B/D – FGFR inhibitors: Infigratinib, Futibatinib, or Pemigatinib combination dose response matrices in the presence of absence of FGF10. 1= 100% viability and 0= 0% viability post indicated treatment. B) Biparatopic antibody B/C – FGFR inhibitors: Infigratinib, Futibatinib, or Pemigatinib combination dose response matrices in the presence of absence of FGF10. 1= 100% viability and 0= 0% viability post indicated treatment. C) Heatmap showing Bliss scores calculated from dose response matrices D-E) Viability of NIH3T3 cells stably expressed FGFR2-ACHYL1 with V565I or V565F mutations treated with bpAb-B/D, bpAb-B/C, or IgG1. F) Immunoblot analysis of NIH3T3 cells stably expressed FGFR2-ACHYL1 with V565I or V565F treated with bpAb-B/D, bpAb-B/C, or IgG1. G-H) Quantification of internalization/degradation signals in FGFR2-ACHYL1 with V565I or V565F expressing NIH3T3 cells treated with biparatopic antibody bpAb-B/C, bpAb- B/D, or IgG1 from 0-38hrs post incubation. I) Viability of CCLP-1 cells stably expressed FGFR2–PHGDH fusion with V565F mutation upon treatment with IgG1, bpAb-B/D or bpAb-B/C alone or in combination with Infigratinib (% compared to IgG1 treated control). J) Immunoblot analysis of CCLP-1 cell line expressing FGFR2-PHGDH with V565F mutation upon treatment with IgG1, bpAb-B/C, bpAb-B/D, IgG1+Infigratinib, bpAb- B/C+Infigratinib, or bpAb-B/D+Infigratinib for 5hrs. K) Deletion mutations derived from 4 different patients and the respective FGFR2 ECD. L) Viability of 4 patient derived N-terminus oncogenic mutants upon treatments with IgG1, bpAb-B/C, or bpAb-B/D as indicated (% viability compared to IgG1). M) Immunoblot of NIH-3T3 cells bearing an FGFR2 H167_N173 in-frame deletion allele (patient 2) after treatment with IgG, bpAb-B/C, bpAb-B/D or the relevant parental antibodies. Data are mean ± SEM. ns=not significant, *P < 0.05, **P < 0.01, ***P < 0.001, ****P < 0.0001 by One-way ANOVA multiple comparisons.

Diverse secondary FGFR2 kinase domain mutations drive clinical resistance to each of each FGFR TKI studied to date (*3, 34, 35*). Given the intracellular location of the kinase domain, we hypothesized that the biparatopic antibodies might remain active against these mutations. To test this hypothesis, we selected the gatekeeper mutations V565I and V565F, which are common mechanisms of resistance to the approved FGFR inhibitors. NIH3T3 cells stably expressed FGFR2-ACHYL1 with a V565I or V565F mutation were resistant to infigratinib (Fig. S7A) but were sensitive to bpAb-B/C and bpAb-B/D, showing inhibition of both growth and downstream signaling (Fig. 7D-F). Moreover, bpAb-B/C or bpAb-B/D induced lysosomal degradation of the FGFR2 fusion in these cells as assayed by anti-Fc Fab fragment conjugated pH-sensitive fluorophore (Fig. 7G, H), similar to what we observed in NIH3T3 cells expressing WT FGFR2 fusions (Fig. 6F-H). Given the complexity of resistance mechanisms in patient tumors, which may implicate multiple oncogenes and bypass mechanisms, we modeled the efficacy of our antibodies in the FGFR1-dependent cholangiocarcinoma cell line, CCLP-1, stably transduced to express the FGFR2-PGHDH WT or FGFR2-PHGDH-V565F alleles (Fig.S7B). CCLP-1 parental cells as well as CCLP-1 cells expressing FGFR2-PHGDH WT were sensitive (IC50<2nM), while FGFR2-PHGDH V565F cells were resistant (IC50>2000 nM) to infigratinib (Fig.S7C). To determine the dose of infigratinib to use in combination studies (in order to suppress the concurrent FGFR1 activity), we determined the infigratinib concentration that sensitized cells expressing FGFR2-PHGDH WT but not FGFR2-PHGDH V565F (0.15uM). Treatment with bpAb-B/C or bpAb-B/D in combination with infigratinib significantly suppressed growth of V565F resistant mutants and re-sensitized the CCLP-1 resistant cells towards infigratinib, indicating robust suppression of the introduced FGFR2 resistance allele (Fig.7I). In addition, co-treatments of infigratinib and bpAb-B/C or bpAb-B/D inhibited FGFR2 signaling (Fig.7J). These results support the use of bpAb-B/C and bpAb-B/D to overcome secondary FGFR2 kinase domain mutations.

In addition to FGFR2 rearrangements, a recent study has revealed that activating in- frame FGFR2 ECD deletions occur in ∼3% of ICC patients. Patients with these FGFR2 ECD deletions responded well to FGFRi treatments, suggesting that these ECD mutations are oncogenic drivers(*36*). Since these mutations are located in the ECD, it is possible that they might lack sensitivity to our biparatopic antibodies. To determine whether bpAb-B/C or bpAb-B/D has activity against oncogenic FGFR2 ECD in-frame deletion mutations, we engineered NIH3T3 cells to stably express 4 patient-derived FGFR2 ECD deletion mutations (Fig.7K). Compared to NIH3T3 cells expressing FGFR2- WT, cells expressing deletion mutations had increased transformation capacities and receptor dimerization as analyzed by soft-agar assay and NanoBiT assays, respectively (Fig.S7D-G). In addition, these mutants had elevated FGFR2 downstream phosphorylation, which was blocked by infigratinib, confirming their FGFR2 dependency (Fig.S7H). While bpAb-B/C or bpAb-B/D had moderate activities against patient 1 and 3- derived mutants, to our surprise, both bpAb-B/C and bpAb-B/D effectively inhibited growth of patient 2 and 4 variants (Fig.7L). These results correlated with the decrease in phosphorylation of FGFR2 downstream signaling for the H167_N173Del (patient 2) variant (Fig.7M). Importantly, levels of FGFR2 decreased upon bpAb-B/C and bpAb-B/D treatments, suggesting that receptor internalization and degradation mediate the observed growth inhibition (Fig.7M). Crucially, mutations found in patients 1-4 are predicted to alter the three-dimensional structure of FGFR2 D2 and D3 domains (*36*) and may consequently affect the binding affinities of bpAb-B/C and bpAb-B/D with D1 and D2 binding arms. Nevertheless, the fact that bpAb-B/C and bpAb-B/D remain effective against patient 2 and 4 variants suggest that as long as the binding avidities of D1 and D2 binders are sufficient to establish intermolecular interaction and trigger internalization, the bpAb-B/C and bpAb-B/D should be effective. These data demonstrate that bpAb-B/C and bpAb-B/D have activities against intracellular kinase domain mutations and specific patient-derived FGFR2 ECD oncogenic deletions. Together with the observed synergy, these data support the notion of combining FGFR1-3 inhibitors with FGFR2 biparatopic antibodies.

## DISCUSSION

In this study, we established that the FGFR2 ECD is required for the oncogenic activity of FGFR2 fusions. A series of monoparatopic antibodies against FGFR2, however, were largely ineffective at blocking downstream signaling. Accordingly, we systematically generated biparatopic antibodies against a diverse combination of epitopes that span three domains on the FGFR2 ECD. Through unbiased phenotypic screening using cancer growth inhibition as a functional readout, we selected two biparatopic antibody candidates that achieved highest efficacy in vitro and confirmed their therapeutic activities in FGFR2 fusion ICC xenograft models in vivo. The antibodies had synergistic combination activity with FGFR2 TKIs and had activity against gatekeeper kinase mutations as well as N-terminal oncogenic FGFR2 alterations in the ECD. Overall, our work highlights the therapeutic potential of these antibodies in ICC and presents a framework for the development of biparatopic antibodies more broadly.

A variety of modes of action of biparatopic antibodies might contribute to their efficacy. Upon binding to its target, the biparatopic antibody could 1) exert agonistic activity by mimicking the ligand-induced receptor activation(*37*), 2) act as a true ligand-antagonist blocking the ligand interaction and downstream signaling activation, or 3) induce receptor internalization and degradation through intermolecular crosslinking and complex formation. Critically, only the latter mode of action can inhibit ligand-independent receptor activation and sustainably downregulate signaling pathway to reduce tumor growth. In this work, we have shown mechanistically that the abilities of bpAb-B/C and B/D to effectively inhibit ligand-independent FGFR2 fusion activation are mediated through enhanced receptor internalization and lysosome-mediated receptor degradation, which results in tumor growth inhibition in vivo.

Recent advances have been made in the field of targeted protein degradation utilizing endo-lysosomal pathways, such as lysosome-targeting chimeras (LYTACs) and antibody-based PROTAC (AbTAC) platforms. Despite their promises at eliminating soluble proteins, the success of these platforms at targeting membrane receptors relies on the endogenous trafficking kinetics of specific RTKs, lysosome targeting receptors, or transmembrane E3 ligases involved as well as their expression and colocalization(*38, 39*). Moreover, such antibodies require further modifications beyond the standard IgG format. Biparatopic antibodies, on the other hand, can be systematically designed against receptors such that the specific epitope combinations can promote receptor binding, trafficking, and degradation of target receptors. If such antibodies can achieve comparable target degradation, they would be accompanied by the advantages of a standard IgG format, including long half-life, high specificity, ability to recruit effector functions, and low immunogenicity(*40*). Thus, the rational engineering and screening of biparatopic antibody platforms may provide a simple yet powerful approach to target a broad range of receptor oncogenes.

Acquired secondary mutations in the FGFR2 kinase domain are an important mechanism of resistance to FGFR TKIs. Although next-generation covalent FGFR TKIs with broader spectrum activity against these mutations have been developed, on-target resistance remains a significant limitation to monotherapy with these agent (*3*). We provide proof-of-concept data that biparatopic antibodies bpAb-B/C and bpAb-B/D targeting the FGFR2 ECD can overcome various kinase domain resistance in FGFR2 fusions. Indeed, previous studies have leveraged antibody or antibody combinations to overcome acquired resistance in other cancer settings, such as in the case of EGFR(*41, 42*). Thus, biparatopic antibodies with high activity and low toxicity have the therapeutic potential to target various forms of RTK resistance to small molecule kinase inhibitors.

We and others have shown that dual inhibition of oncogenes using two targeted agents having non-overlapping patterns of cross-resistance can delay or prevent the occurrence of on-target resistance(*43, 44*). Specifically, dual targeting of BCR-ABL oncogene with a combination of allosteric and catalytic ABL inhibitors acting at distinct sites are non-cross resistant and eradicate CML tumors in preclinical models(*44*). Similarly, based on the observed synergy between bpAb-B/C and bpAb-B/D and FGFR inhibitors (Fig.7) we speculate that combination treatments of FGFR2 biparatopic antibodies and pan-FGFR inhibitors might delay or prevent the emergence of acquired resistance. A considerable advantage of highly active antibodies is the relative ease of combining such agents with small molecule inhibitors, as it has often been difficult to create well-tolerated combinations of targeted agents.

In all, our work has uncovered potent FGFR2 biparatopic antibodies as potential targeted treatment for FGFR2-driven ICC. Our results demonstrated that the engineering of biparatopic antibodies has the potential to lead to more effective and targeted treatments for a wide range of cancers.

## METHODS

### Generation of DNA constructs and cell lines

FGFR2-ACHYL1(*2*), FGFR2-BICC1(*2*), and FGFR2-PHGDH(*3*) sequences were previously described as referenced. FGFR2-ACHYL1 and FGFR2-BICC1 constructs were synthesized and cloned into MSCV vector. FGFR2 ECD with Ig subdomain deletions were generated based on FGFR2-BICC full-length sequence without Ig1 (D1), Ig2 (D2), and in Ig2-3 (D2+3). All construct maps were sequence validated and aligned using SnapGene software.

For FGFR2 ECD WT and mutants, NIH3T3 and HEK-293T cells were transiently transfected with FGFR2-BICC1 or its variants and were selected with 2ug/mL puromycin for 3 days prior to performing assays. 6 parental antibodies and anti-human IgG1-FITC (Jackson Lab) were used as primary and secondary antibodies respectively to validate the Ig-specific deletion mutants. Analysis was done using FlowJo v.10.8 software. ICC13-7 and CCLP-1 cholangiocarcinoma patient-derived cell lines were gifts from the Bardeesy lab (N.B.) and were authenticated via STR profiling.

### Biparatopic antibodies design and generation

To generate biparatopic antibodies, controlled Fab arm exchange reactions were performed where F405L and K409R containing antibodies were mixed according to the protocol(*24*). To assess the quality and concentration of the bispecific antibodies, SDS- PAGE, SEC-HPLC and Mass Spectrometry analysis were performed.

### Dimerization Assay

For NanoBiT constructs, FGFR2-WT, FGFR2-ACHYL1 and FGFR2-BICC1 were C- terminally tagged with Small BiT or Large BiT derived from NanoLuc (Promega). HEK293T cells were stably or transient transfected. Nanoluc substrate (Nano-Glo® Live Cell, Promega) was added the mixture according to the manufacturer protocol. The luciferase activity was measured by EnVision plate reader (PerkinElmer).

### Immunohistochemistry

Tumors were surgically removed and placed in 10% neutral buffered formalin for 24 hours and followed by 70% ethanol until paraffin embedded. Hematoxylin solutions were used for counterstaining.

### Transformation Assays

*Focus formation assay*: NIH3T3 stably expressing FGFR2 fusions were plated in 6-well plate in triplicate. Cells were grown for 7-10 days, plates were imaged, and the number of foci were blindly counted. *Soft agar colony formation assays*: NIH3T3 cells stably expressing patient-derived oncogenic FGFR2 variants were plated in 6 well-plates with 0.5% Select Agar. Cells were cultured for 2-3 weeks, and colonies were imaged, and colony numbers were determined using ImageJ and Prism software.

*BaF3 transformation assay*: BaF3 cells were resuspended in RPMI media + 10%FBS. Cells were seeded in 6-well plate and were split every 3 days. For each split, Cell-titer Glo was used to measure the cell viability compared to original seeding density and the new seeding density was determined. Cumulative population doublings were calculated over the period of 15-20 days.

### Binding affinity and epitope binning assays

#### MSD-SET (Meso Scale Discovery-Solution Equilibrium Titration)

Measurements were performed according to the previously published protocol(*45*). The MSD SET experiments were carried out in 96-well from MSD (PN: L15XA-3) using assay buffer PBS 1x pH7.4, 0.1% BSA (Sigma-Aldrich) w/v, 0.02% P20 (ThermoFischer). Experiments were performed as independent duplicates.

#### BLI-Octet (Bio-Layer Interferometry)

Binding kinetics (ka, kd) and affinity (KD) were measured in an Octet system RED96e. Antibodies were captured by Anti-Human Fc capture biosensor (AHC). hFGFR2 ECD was used as an analyte, association and dissociation of the analyte to the captured antibody was monitored for 600s. Data were analyzed using the Octet Data Analysis software HT 12.0. Sensorgrams were fitted to a 1:1 binding model where kinetic rate Ka and Kd were globally fitted.

#### Epitope binning

Epitope binning experiments were performed in an Octet system RED96e. To perform an in tandem epitope binning experiment, hFGFR2 ECD was captured on streptavidin sensor (SA). The cycle starts with the capturing of biotinylated ligand followed by a “primary” antibody (Ab1) binding step where Ab1 interaction is monitored for 600s. Data were blindly analyzed using the Octet Data Analysis software HT 12.0 and R Studio “pvclust” according to Octet Application note n.16.

#### Flow cytometry

*Antibody affinity assay*: NIH3T3 cells expressing full-length and FGFR2-BICC1 variants (D1, D2, D3, or D2+D3 deletion variants), SNU-16 cells, or parental BaF3 cells (neg control) per tube were incubated with parental antibody A-F (NIH3T3) or at serial dilutions of 0-10mg/mL (SNU-16) in 1xPBS (*46*). Cells were washed three times with FACS buffer (1xPBS, 1% BSA, 5% FBS) and incubated with anti-human IgG Alexa Fluor 488 secondary antibody, washed, and analyzed on a SA3800 Spectral Analyzer (Sony Biotechnology). Data were analyzed using FlowJo® v.10 software and fit in GraphPad Prism 9 using a ligand-binding quadratic equation to obtain Kd values.

#### Fabfluor receptor degradation

NIH3T3 or ICC13-7 cells were seeded in 96 well-plate (Corning, Catalog#3595). Incucyte® Fabfluor-pH Antibody Label reagents (*31*) stock concentration at 0.5mg/mL were mixed and incubated with each antibody according to the manufacturer’s protocol. Analysis was done using Incucyte Basic Analyzer and Total Integrated Intensity Per Well (RCU/OCU x µm^2^ /Well) was quantified as a readout using Incucyte software v2019B.

#### Growth inhibition Assay

Engineered BaF3 cells expressing FGFR2-PHGDH and FGFR2-ACHYL1 cells were seeded in RPMI + 10%FBS media in 96 well-plates (Corning, Catalog#3904). Parental antibodies, biparatopic antibodies, or IgG1 control were added at 15 serial concentrations ranging from 0 to 1uM in the presence or absence of FGF10. Viability was determined using CellTiter-Glo™ 2.0 (Promega) at day 5 post treatment according to the manufacturer’s instructions.

#### Mouse xenografts assay

BaF3 cells expressing FGFR2-PHGDH or ICC13-7 cells in a total volume of 200uL were subcutaneous implanted in the right flank of 7–9-week-old female BALB/c scid mice. At a tumor size of ∼250mm^3^ (BaF3) or ∼150mm^3^ (ICC13-7), mice were randomized into 10 groups, 10 mice per treatment group. Biparatopic antibodies, parental antibodies, or IgG1 were IV administered twice per week and tumor sizes were measured by caliper every 3- 4 days for 25 days (BaF3) and 38 days (ICC13-7). Tumor volume was calculated by the modified ellipsoidal formula: V = 0.523 x (L x W^2^) where L = the greatest longitudinal diameter and W = the greatest transverse diameter (width). All experiments were conducted under protocol 0121-09-16-1 approved by the Broad Institute’s Institutional Animal Care and Use Committee (IACUC).

#### ELISA assay

Blood samples were collected from the submandibular veins of mice at 1, 24, and 72 hours post the last dose of the treatment before the harvest. Levels of plasma antibody were measured with the Human IgG Total ELISA Kit (Sigma Aldrich) per manufacturer’s instructions. The absorbance was measured with EnVision (PerkinElmer).

#### Phospho-receptor tyrosine kinase profiling

Protein was prepared per protocol (R&D Systems): Cells were harvested in lysis buffer provided in kit with protease and phosphatase inhibitors added before use. Membranes were exposed to X-ray film (Fuji) for multiple exposure times and dots were mapped using reference spots provided and analyzed for relative intensity using ImageJ.

## Acknowledgements

We thank the members of the Flow Cytometry Core Facility at the Broad Institute for cell analysis and the Comparative Medicine Department at the Broad institute for animal care.

## Author contributions

S.C. and W.R.S. conceived, designed, analyzed the experiments, and wrote the manuscript. S.C., S.O., D.T.F., J.K., F.P.R., T.Y.S., and M.C. performed antibody validations, in vitro activity assays, and mechanistic validation experiments. A.S., S.C. characterized antibody binding epitopes and affinities. J.K., S.C., D.J.R., L.C., performed in vivo xenografts experiments. D.K. provided antibody-antigen structural insights. D.T.F., J.K., A.A., R.D., Y.Y.T., Y.H., processed and analyzed xenograft-derived samples. N.B. provided ICC models and critical insights into ICC biology and FGFR inhibitors. All authors reviewed and edited the manuscript.

